# Gene Expression Variability with Feedback Regulation Implemented via Protein-Dependent Cell Growth

**DOI:** 10.64898/2026.04.13.718123

**Authors:** Iryna Zabaikina, Pavol Bokes, Abhyudai Singh

## Abstract

Variability in gene expression among single cells and growing cell populations can arise from the stochastic nature of protein synthesis, which often occurs in random bursts. This study investigates the variability in the expression of a growth-sustaining protein, whose concentration is regulated by a negative feedback loop due to cell growth-induced dilution. We model the distribution of protein concentration using a Chapman-Kolmogorov equation for single cells and a population balance equation for growing cell populations. For single cells, we derive an explicit solution for the protein concentration distribution in state space and represent it as a Bessel function in Laplace space. For growing populations, we find that the distribution satisfies a Heun differential equation with singular boundary conditions. By addressing the central connection problem for the Heun equation, we quantify the population-level protein distribution and determine the Mathusian parameter, which characterizes population growth. This work provides a comprehensive analytical framework for understanding how stochastic protein synthesis impacts gene expression variability and population dynamics.

## I. Introduction

Gene expression, the process by which genetic information is used to produce functional molecules such as RNA and proteins, is a fundamental driver of all cellular functions, including growth, division, and adaptive responses to environmental changes. This process is inherently stochastic, leading to significant cell-to-cell variability in not only protein concentration but also cell cycle progression, metabolic regulation, and overall growth dynamics. As a result, cells employ regulatory mechanisms, such as feedback loops, to modulate gene expression.

We focus on proteins that actively promote cell growth, such as (i) growth factors [1], [2], which activate signalling pathways to stimulate cell proliferation and survival; (ii) *Myc* transcription factors [3], [4], which regulate metabolic pathways and drive cell-cycle progression; (iii) cell growth-activating enzymes [5], [6], [7], which can regulate cell-cycle progression and checkpoint control. Such proteins are essential for normal cell-cycle regulation. Their dysregulation has been linked to hyperproliferation in cancer cells [8] and to oncogenic transformation [9], [10].

At the single-cell level, these proteins form a negative feedback loop: dilution due to cell expansion lowers protein concentration and thereby slows further growth. This mechanism promotes homeostasis by limiting excessive fluctuations in protein levels and has been observed experimentally in both bacterial and mammalian systems [11]. It may also lead to deviations from classical exponential growth, yielding a hearly linear growth pattern [12], [13]. At the population level, because cell division is closely tied to cell growth and changes in cell volume [14], [15], this growth-control mechanism can substantially affect proliferation dynamics and lead to discrepancies between single-cell and population-level protein distributions [16]. Complementary positive-feedback scenarios have also been investigated in related stochastic models of gene expression in proliferating cells [17], [18]. These observations motivate a refined mathematical framework for studying the phenomenon at both singlecell and population levels.

To study protein concentration variability under protein-dependent cell growth, we adopt continuous-valued frameworks. At the single-cell level, we use the widely applied piecewise-deterministic Markov process (PDMP) framework [19], which captures cellular dynamics with both deterministic components (e.g., degradation or dilution) and stochastic components (e.g., protein synthesis or cell division). At the population level, instead of aggregating individual PDMP trajectories, we construct a separate model based on the population balance equation (PBE), a standard framework for describing the distribution of cell states in a growing and dividing population [20], [21], [22]. Stationary protein distributions are then obtained by combining the PBE with branching process theory accounting for birth and death events in the population [23], [24]. Due to the complexity of the PBE, its solutions, including protein distributions, often involve special functions [25], [26], [27]. In particular, the Heun function appears in several protein distribution models [28], [29], [30], [31] and also in this work. These equations present substantial analytical and numerical challenges, which we address in this work.

We develop a mathematical framework that describes protein-dependent cell growth at both single-cell and population levels. We solve a version of the PBE with negative feedback, combining analytical and numerical techniques. Our approach extends existing models by explicitly incorporating a negative feedback mechanism, allowing a deeper understanding of its effect on protein distributions.

## II. Negative feedback: single cell

We start with a single-cell model, in which the protein concentration increases during short, intense periods of synthesis known as *bursts*. These bursts occur randomly in time according to a Poisson process with rate *α*. The size of each burst follows an exponential distribution with mean *β* [19]. Note that bursts may represent either transcriptional or translational events, which we combine into a single effective burst. This concentration-based model relies on the experimentally observed scaling of gene product numbers with cell size [32]; hence, we work with protein concentration, whose time evolution is independent of cell size. We assume that the protein is relatively stable, so that between bursts its concentration decreases only through cell growth-induced dilution. The dilution rate is characterized by *γ*(*x*) [33], [34], and thus the deterministic dynamics are governed by the ordinary differential equation

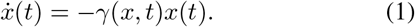

Since the protein promotes cell growth, higher protein concentrations increase the dilution rate, thereby creating an effective negative feedback loop (Fig. 1A). We model this dependence using the Michaelis–Menten response function:

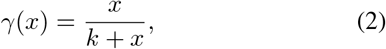

where *k* is the half-saturation constant. In (2), the maximum growth rate is set to one without loss of generality.

**Fig. 1.**
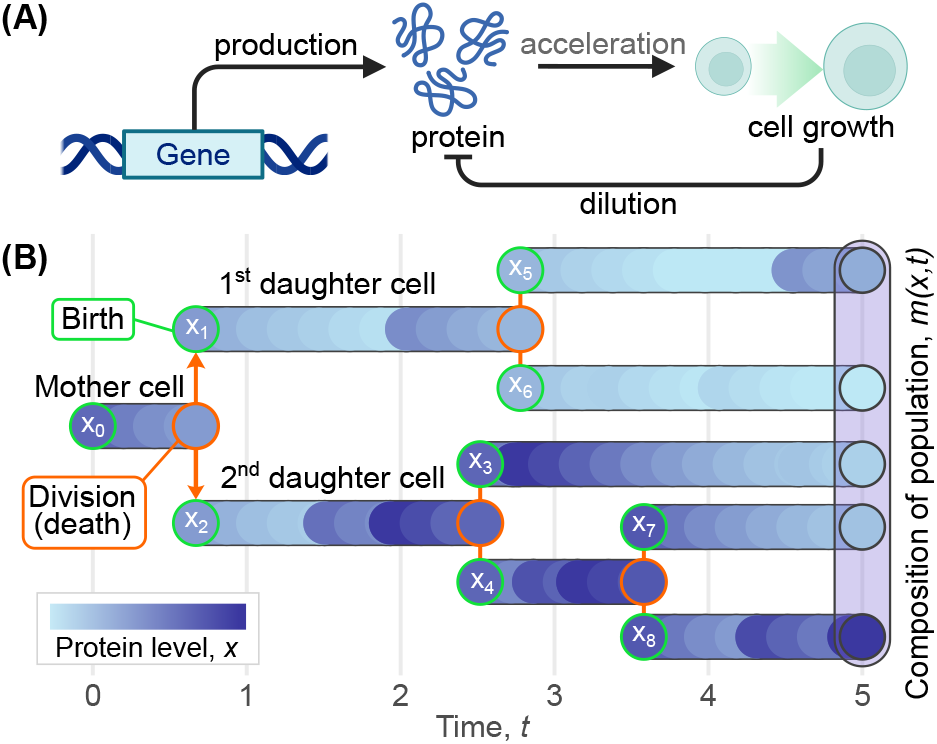
(A) Scheme of the process at the single-cell level. The protein accelerates cell growth, thereby creating negative feedback through increased dilution. (B) Representation of the sample population as a lineage tree. Each branch corresponds to one independent cell and describes how its protein concentration changes over time. The colour intensity is proportional to the protein concentration. Each new cell is assigned an index *i*, and *x*_*i*_(*t*) denotes the protein concentration in the *i*th cell at time *t* (in the figure, it is shown at the moment of birth). The empirical population density *m*(*x, t*) is determined by all cells alive at time *t*.

The absence of feedback, *k* = 0, corresponds to unregulated dilution (the growth rate does not depend on protein level). In this case, the stationary protein concentration follows a gamma distribution with shape *α* and scale *β* [35]. Cell volume then grows exponentially and protein concentration decays exponentially between burst events. The inclusion of the half-saturation constant (*k >* 0) determines how sensitively cell growth, and hence dilution, responds to protein concentration. When the protein concentration is low (*x < k*), growth is slow, and the dilution rate remains low. For sufficiently high protein concentrations (*x > k*), growth accelerates, causing faster dilution. Thus, larger *k* corresponds to lower sensitivity, meaning that higher protein concentrations are required to achieve a given growth rate.

The time evolution of the probability density *p*_sc_(*x, t*) in a single cell is described by the Chapman-Kolmogorov equation [35], [36]:

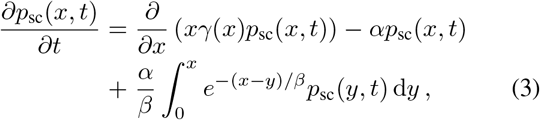

where *y* is the integration variable representing possible pre-burst protein concentrations.

To obtain the steady-state distribution, we set the time derivative to zero and introduce an auxiliary function 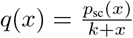. We convert the steady-state equation into an integral equation in *q*(*x*) of a convolution type. Subsequently, we take the Laplace transform *Q*(*s*) = ℒ_*s*_ *q*(*x*) and obtain a linear second-order differential equation:

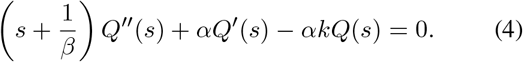

After a shift in the Laplace variable, (4) becomes the modified Bessel differential equation [37]; the general real-valued solution is a linear combination of the modified Bessel functions of the first and second kinds, *I*_1−*α*_(·) and *K*_1−*α*_(·), respectively. Since *Q*(*s*) must remain bounded, and *I*_1−*α*_(·) is unbounded, the solution will contain only the branch involving *K*_1−*α*_(*x*).

Using the integral representation of *K*_1−*α*_(*x*) [38], we obtain the following explicit form of the steady-state probability density function for the protein concentration with negative feedback on dilution:

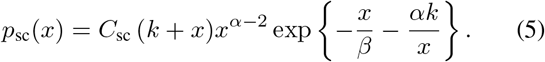

The value of the constant *C*_sc_ is chosen so that *p*_sc_(*x*) integrates to one:

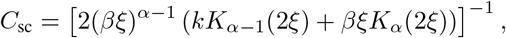

where 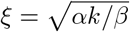. Note that

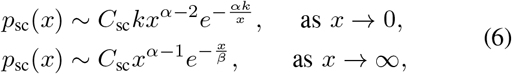

i.e. *p*_sc_ is exponentially small near *x* = 0 and *x* = ∞; consequently, all polynomial moments of *p*_sc_ are finite.

The *n*-th protein moment in the single-cell model is

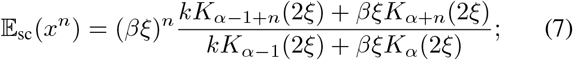

here 𝔼(·) denotes the expected value, namely 𝔼_sc_(*x*^*n*^) = ∫ *x*^*n*^*p*_sc_(*x*) d*x*.

We analyse the effect of the feedback from two perspectives. First, we fix the production parameters *α* and *β*, and hence the average production rate *αβ* (Fig. 2A). We observe that the probability mass near low protein concentrations decreases. This is because the left tail of *p*_sc_(*x*) is governed by the exponential factor 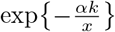, which becomes steeper for higher *k*. A higher half-saturation constant requires a higher protein concentration before dilution accelerates, so protein accumulates longer, widening the distribution and shifting its mode to the right. Note, that for any *k* the right tail is governed by the exponential term 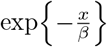, so dilution always remains effective enough to counterbalance production, and the distribution always exists for all admissible parameter values. The following approximation for the protein mean holds once the feedback is strong enough:

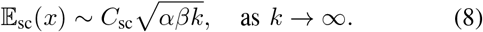

**Fig. 2.**
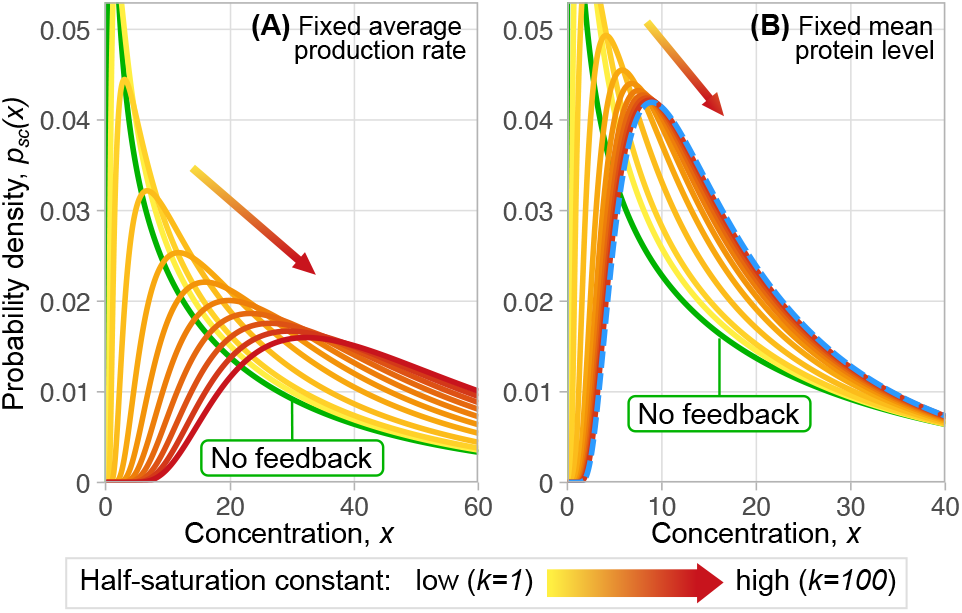
The protein distribution in the single-cell model, given that: (A) the average production rate is fixed (*αβ* = 25, *β* = 40); (B) the mean protein concentration is fixed (𝔼_sc_(*x*) = 25), achieved by adjusting *α* for each value of *k* according to (7). Separately, we highlight the case *k* = 100 by the dashed blue line. In both panels, the green line corresponds to the case of no feedback (*k* = 0), and it is identical in panels (A) and (B). The half-saturation constant *k* is shown as a gradient from yellow to red, with larger *k* shifting the threshold of fast dilution to higher protein concentrations.

Second, we fix the mean protein concentration 𝔼_sc_(*x*) (Fig. 2B); the burst frequency *α* is then decreased as per (7) as *k* increases. Namely, (8) implies that 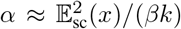 for sufficiently large *k*. Then the limiting distribution 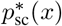 asymptotically behaves as follows:

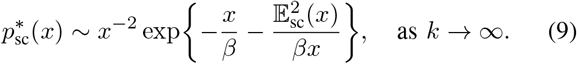

As we discussed, larger values of *k* shift the point at which dilution accelerates to higher concentrations, but the cell compensates by lowering the burst rate, thereby maintaining homeostasis. Further increasing *k* does not alter the single-cell concentration distribution.

## III. Negative feedback: cell population

We now extend our analysis from the single-cell level to a population framework. We start with the cell cycle of an individual cell that is characterized by the birth time *t*_*b*_ and the division time *t*_*e*_. During this period, the protein dynamics in each individual cell in the population has the dynamics as in Section II. The proteins of interest (e.g., growth factors) affect the cell cycle as follows: they accelerate cell growth, while larger cell size triggers specific checkpoints in the cell cycle, and the division event occurs. Hence, higher protein concentrations accelerate the rate of cell division. To capture this effect, we assume that the rate at which the cell cycle ends (and the cell divides) depends on the current protein concentration through the same response function *γ*(*x*) that governs dilution. Thus, the probability that the cell cycle ends in the interval [*t, t* + *dt*] is given by:

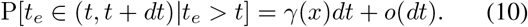

The completion of the cell cycle triggers a branching (division) event: the current process (the mother cell) is terminated and replaced with two new processes (daughter cells), each of which inherits a half of the mother’s protein content and a half of the mother’s cell volume. Hence the concentrations (amount of protein per unit volume) remain the same through the division event.

The daughter processes evolve then independently of each other, and we continue to track both of them. The sample trajectory and corresponding lineage tree are shown in Fig. 1. The population net growth rate then represents the cumulative effect of all birth-death processes. We assume that the net growth rate is proportional to the current density of all cells at a given concentration *x*.

We index cells in the population by *i* (as shown in Fig. 1). For each cell *i* in the population, *x*_*i*_(*t*) denotes its stochastic protein concentration trajectory during its lifetime, from the birth time 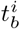 to the division time 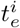. The composition of the population at time *t* can be represented by the empirical population density:

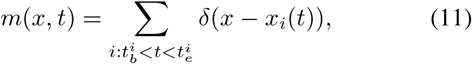

where the sum runs over all cells alive at time *t*. The measure {*m*(*x, t*)} is empirical because it aggregates a finite set of observations *x*_*i*_(*t*) and is random because these observations are realizations of a stochastic process. It is not normalized: ∫*m*(*x, t*)*dx* = *N*(*t*) is the number of living cells at time *t*. Then *m*(*x, t*) can be viewed as a measure-valued Markov process since the future dynamics of the population depend only on the present state of the measure *m*(*x, t*).

Let the population density function *h*(*x, t*) be the average number of cells with protein concentration *x* at time *t*:

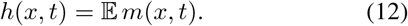

We assume that *h*(*x, t*) is a sufficiently smooth function, allowing differentiation and integration as many times as required. Then *h*(*x, t*) satisfies the population balance equation (PBE) [39]:

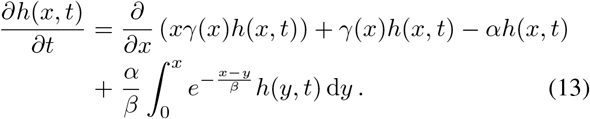

We note that the population balance equation (13) differs from the Chapman–Kolmogorov equation (3) by including the net population growth rate term *γ*(*x*)*h*(*x, t*).

The large-time behaviour of (13) is characterised by the principal eigenvalue *λ*, associated eigenfunction *p*(*x*), and adjoint eigenfunction *u*(*x*). These characteristics can be found using the spectral decomposition:

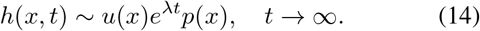

The adjoint eigenfunction is chosen so that it satisfies the biorthogonality condition 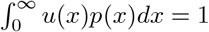. The adjoint eigenfunction determines the dependence of the exponential prefactor on the initial condition so it does not influence the shape of the distribution; thus, it will not be calculated in this work. In the case of the Chapman–Kolmogorov equation (3), the principal eigenfunction *p*(*x*) is the stationary probability density function (5), the principal eigenvalue is *λ* = 0, and the adjoint eigenfunction is trivial, *u*(*x*) ≡ 1.

It follows from (14) that the total population increases exponentially for large times. The rate of the exponential growth, i.e. the Malthusian parameter, is given by the principal eigenvalue *λ*. The normalized empirical population density *m*(*x, t*)*/N*(*t*) converges as *t* → ∞ to the principal eigenfunction of the population balance equation.

Substituting (14) into (13), we obtain an integro-differential equation for the stationary distribution:

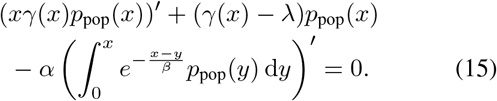

Note that multiplying (15) by *x* and integrating over *x* ∈ (0, ∞) yields

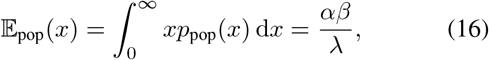

expressing the population mean in terms of the (unknown) Malthusian rate *λ*.

We transform this integro-differential equation to a purely differential one. We carry out the differentiation of the integral term, multiply (15) by *β*, and differentiate the whole equation with respect to *x*; then we add the resulting equation to the original one (15). This transforms (15) into a second-order linear ODE. In terms of an auxiliary function:

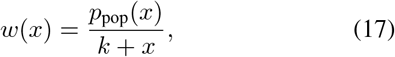

equation (15) becomes the doubly-confluent Heun equation (DCHE) [40]:

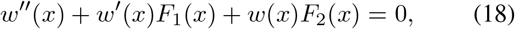

with

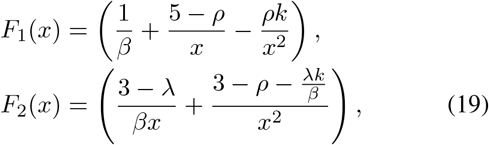

and *ρ* = *α* + *λ*.

Our task is to find the eigenvalue *λ* for which a solution *w*(*x*) to (18) is well behaved (in a suitable sense). This is called a *central connection problem* [41] for the Heun equation, which can be solved numerically.

Equation (18) has irregular singularities of rank one at *x* = 0 and *x* =∞. We use the WKB method [42] to analyze the general solution of (18) at these points, which yields the following asymptotics (details are provided in Appendix II):

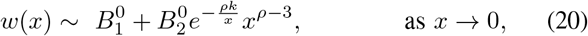

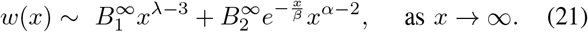

We seek a solution that is exponentially small at both singular points *x* = 0 and *x* =∞, as was observed in the single-cell case (6), i.e. we require that 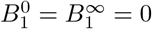.

By numerically integrating an initial-value problem, we construct a solution *w*_*L*_(*x*) to (18) satisfying 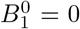. This solution branch is exponentially small as *x* → 0. However, in general, this solution has 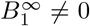, and so does not remain exponentially small as *x* → ∞. Similarly, we construct a solution *w*_*R*_(*x*) to (18) satisfying 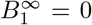, i.e. the branch that is exponentially small as *x* → ∞, but in general 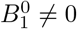.

To obtain a solution that is exponentially small near both singularities, we require that *w*_*L*_(*x*) and *w*_*R*_(*x*) be linearly dependent, which can be tested by the Wronskian

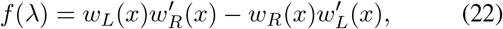

where *x* ∈ (0, ∞) is a suitably chosen test value of the independent variable. The value of *λ*^∗^ for which *f*(*λ*^∗^) = 0 gives the sought-after eigenvalue, and *w*_*L*_(*x*) (or *w*_*R*_(*x*), which is a constant multiple thereof) gives the associated eigenfunction (as shown in Fig. 3).

**Fig. 3.**
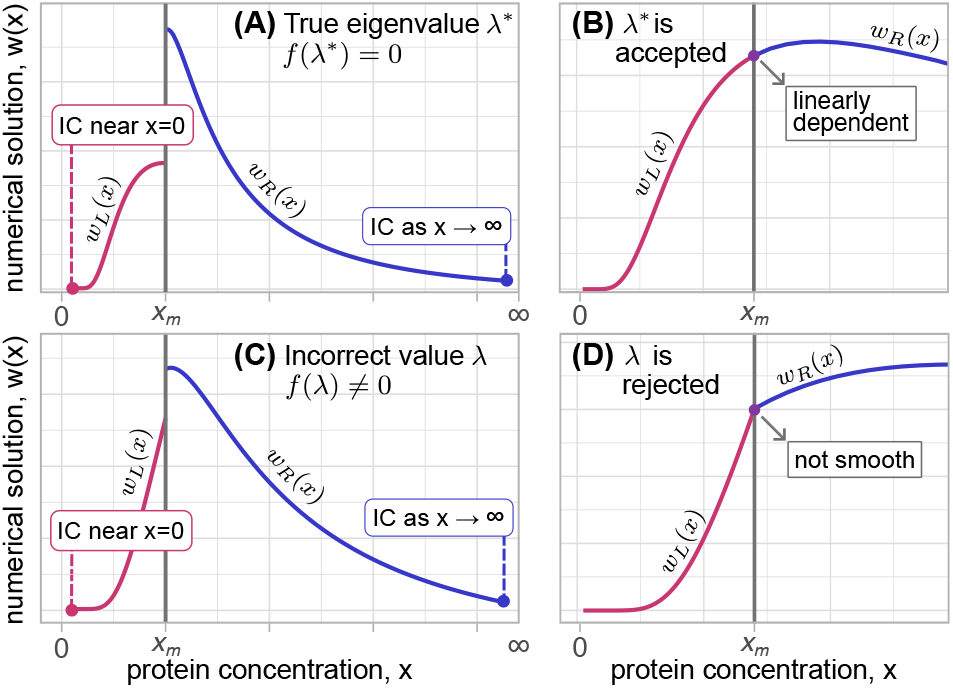
Illustration of the central connection problem. The equation (18) is solved numerically in two parts: two local solutions are computed on the left and right of the matching point *x*_*m*_ by solving initial-value problems started sufficiently close to *x* = 0 and at sufficiently large value of *x*, respectively. The required initial conditions, denoted by IC in the figure, for *x* ≈ 0 and *x* → ∞ are obtained from (20) and (21), respectively. (A, B) For the true eigenvalue *λ*^∗^, we have *f* (*λ*^∗^) = 0, so the two branches match at the meeting point *x*_*m*_ and form a global numerical solution of (18). (C, D) For an incorrect value of *λ*, we have *f* (*λ*) ≠ 0, so the two branches cannot be matched smoothly at *x*_*m*_ and therefore do not form such a solution.

## IV. Results

We analyze how negative feedback on dilution influences protein concentration at both the single-cell and population levels. At the single-cell level, we model protein dynamics using a piecewise-deterministic Markov process. By solving the corresponding Chapman-Kolmogorov equation, we obtain an explicit solution for the steady-state probability density function (5) and the mean protein concentration (7). When the production parameters are fixed, increasing *k* shifts the single-cell protein concentration distribution toward higher values and broadens it (Fig. 2A). However, when we choose *α* so that the single-cell mean 𝔼_sc_(*x*) remains fixed for each value of *k*, this allows us to isolate the effect of negative feedback, and we observe that *p*_sc_(*x*) changes only moderately: it becomes more concentrated, while for sufficiently large *k* it approaches a limiting shape (Fig. 2B).

At the population level, we formulate the population balance equation (13) to describe protein concentrations in the stationary cell population. The ordinary differential equation resulting from (13) is classified as a Heun equation, which arises due to the nonlinearity introduced by feedback and cell proliferation dynamics. All transformations we applied to (13), as well as to its Laplace transform, led either to the general Heun equation or to one of its confluent forms. Its structure with multiple singularities makes finding closed-form solutions rarely feasible. The parameter relations in (18) make some known techniques for reducing DCHE to more common types of equations inapplicable [40], [43], [44]. Thus, we focus on the doubly confluent Heun equation (18), which we solve numerically using a matching technique for inner and outer solutions (Fig. 3). To support the numerical method, we use simulations to guide the choice of numerical parameters (Appendix I) and the WKB expansion to derive the required initial conditions (Appendix II). This approach is expected to be applicable to a broad class of doubly confluent Heun equations, provided that numerical parameters are chosen carefully to maintain solution stability.

To compare the population-level and single-cell distributions, we fix the parameters so that the mean protein concentration in the single-cell model, 𝔼_sc_(*x*), remains fixed. We observe that *p*_pop_(*x*) changes only moderately as *k* increases (Fig. 4A). Moreover, for large *k, p*_pop_(*x*) remains close to *p*_sc_(*x*), although the population-level distribution is slightly shifted to the right (Fig. 4B). The mean protein concentration at the population level, 𝔼_pop_(*x*), approaches a finite limit (Fig. 4C), it shows that growth-coupled negative feedback limits excessive concentration buildup in the cell population. In contrast, in the case of positive feedback on dilution that we studied previously [33], the population distribution becomes nearly uniform, leading to an unbounded increase in mean protein concentration. At the same time, the population level noise remains lower than the single-cell noise and decreases as *k* increases (Fig. 4D), showing that growth-coupled negative feedback also reduces variability at the population level.

**Fig. 4.**
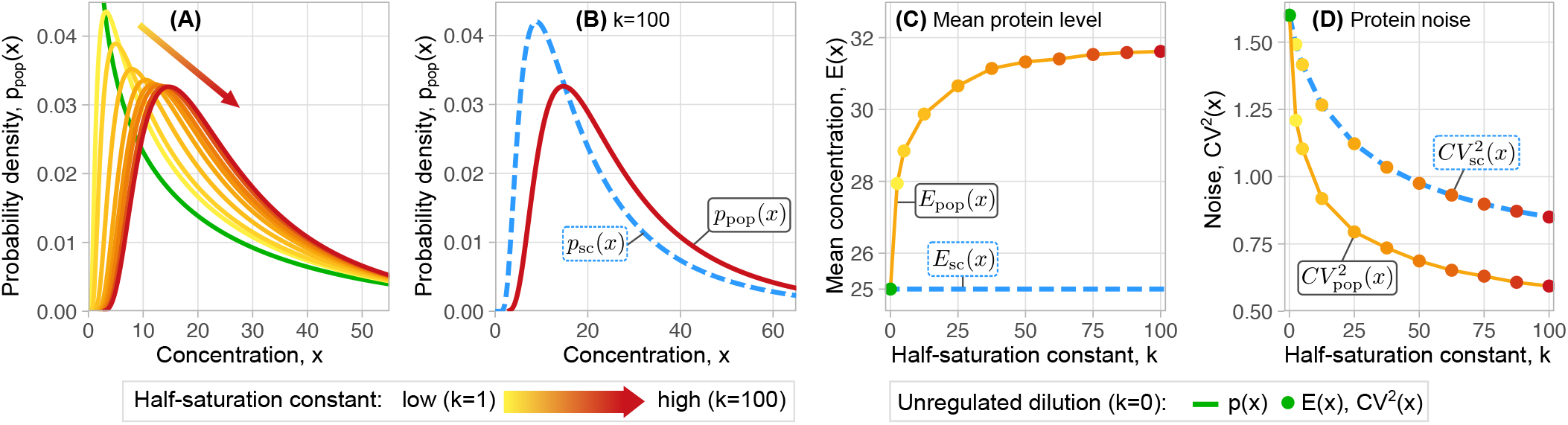
Negative feedback on dilution constrains protein fluctuations within the population. In (A)–(D) parameters are as follows: *β* = 40, *α* is chosen so that 𝔼_sc_(*x*) = 25 for a given value of *k*, as per (7). (A) When *k* is low (*k* = 1, yellow line), the protein distribution resembles that of unregulated dilution (*k* = 0, green line). As *k* increases (lines become more red), left tail becomes lighter and the distribution becomes narrower. (B) Comparison of the *p*_sc_(*x*) (blue dashed line) and *p*_pop_(*x*) (dark red line) for *k* = 4 𝔼_sc_(*x*) = 100. Note that the right tail of *p*_pop_(*x*) is slightly heavier, but overall both distributions are close. (C) For low *k*, the mean protein concentration in the cell population is close to that of unregulated case, but increases rapidly as *k* increases, then growth slows down indicating that 𝔼_pop_(*x*) is a bounded function. (D) Population distribution is less noisy than the single-cell distribution.

From a biological point of view, negative feedback creates a counterbalancing mechanism at both the single-cell and population levels. At the single-cell level, higher protein concentration accelerates cell growth, which increases dilution and counteracts further protein accumulation. At the population level, cells with higher protein levels proliferate faster, but they also dilute the protein more rapidly. Together, these two effects stabilize protein concentration in the growing population and keep the population-level distribution close to the corresponding single-cell one. These findings emphasize the importance of feedback control in cellular systems and provide a foundation for further theoretical and experimental investigations.

## Appendix I

### Simulations

In this appendix, we provide details for the population simulation. We use a recursive algorithm in which a process calls a copy of itself after the condition for cell division is reached. Then the mother cell (process “single cell”) after division splits into two daughter cells (calls two identical processes “single cell”). The whole procedure is described in Algorithm 1.

The deterministic decay of the protein is governed by the ODE *x*^*′*^ = − *xγ*(*x*) (*γ*(*x*) is given by (2)), the solution of which is:

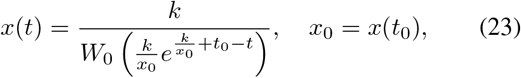

where *t*_0_ is the time of the last event (a burst or a cell division) and *W*_0_(·) is the principal branch of the Lambert *W* function.

Next, as discussed in Section III, the division rate is protein-dependent and given by *γ*(*x*), then the division events arrive as per non-homogeneous Poisson process. Subsequently, the waiting time until the next burst *t*_*e*_ is given by the probability:

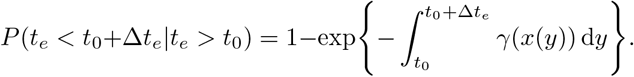

To sample from this distribution, we use the inverse transform technique: then to find Δ*t*, we draw *u*_*d*_ from a standard uniform distribution and evaluate the integral:

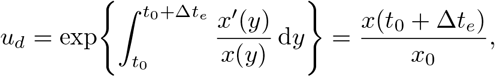

which yields:

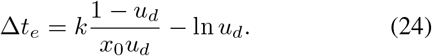

In Section II, we state that the bursts arrive at the constant frequency *α*, then the waiting time until the next burst Δ*t*_*α*_ is an exponentially distributed random variable with mean 1*/α*; the burst size *b* is also drawn from the exponential distribution with mean *β*. We sample both values again using the inverse transform technique:

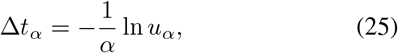

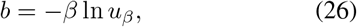

where *u*_*α*_ and *u*_*β*_ are drawn from the standard uniform distribution.

## Appendix II

### The WKB-approximation as *x* → ∞ and at *x* = 0

The numerical method requires local solutions of (18) near its irregular singularities. To obtain it, we perform the WKB-approximation of *w*(*x*), which is based on the assumption that the solution has an exponential form.

First we consider the limit *x* → ∞. To we let *z* = *εx*, so that (18) becomes:

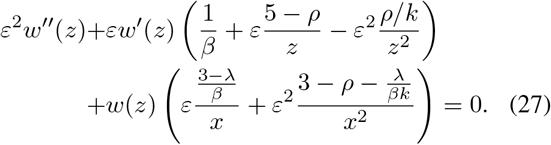

#### Algorithm 1

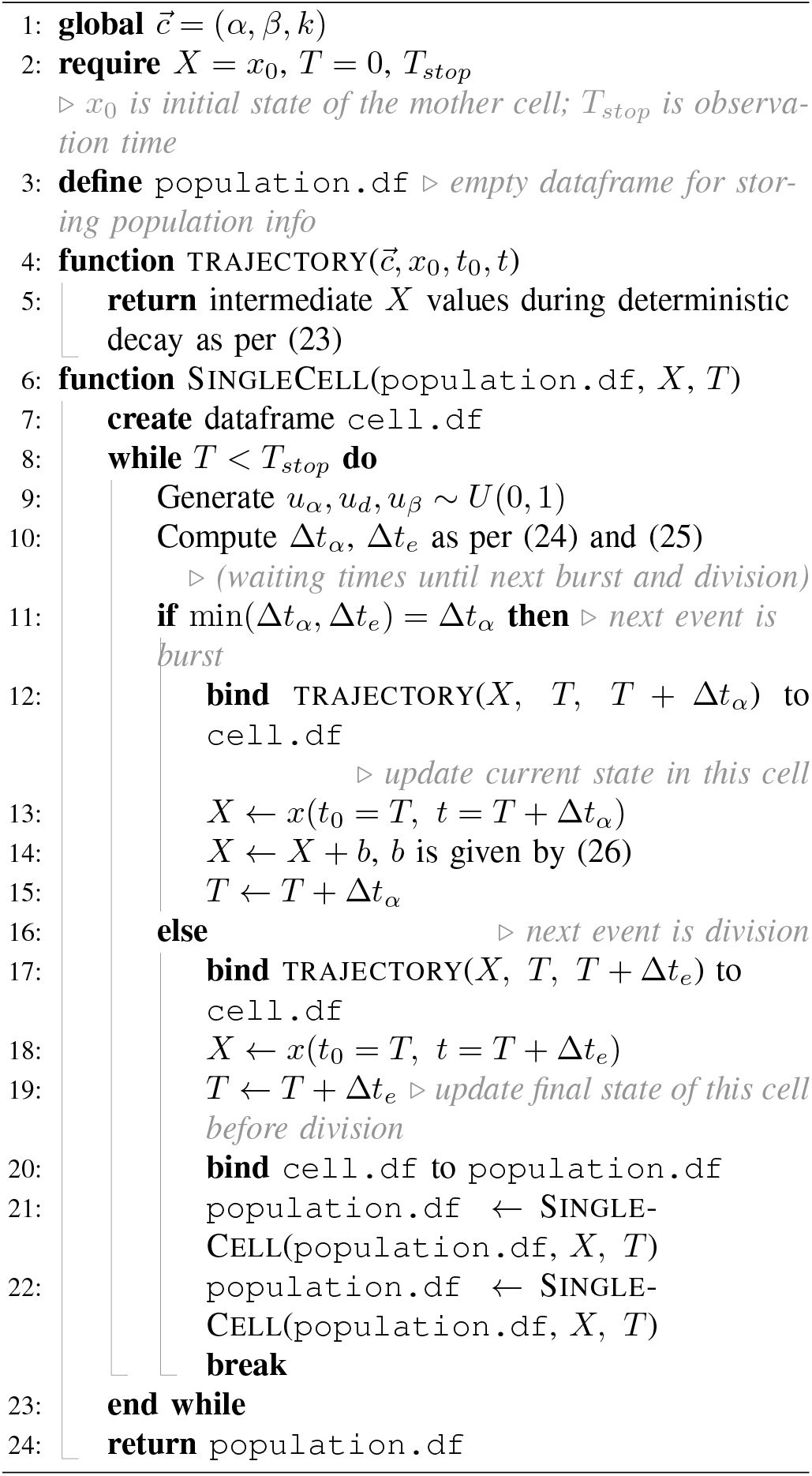

We are looking for the asymptotic expansion of the form:

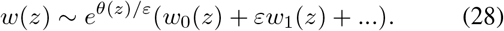

Substituting it into the ODE yields a system of equations, which determines *θ*(*z*) and *w*_0_(*z*):

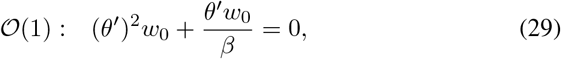

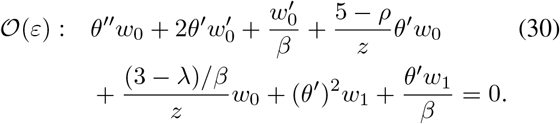

The eikonal equation (29) has two solutions:

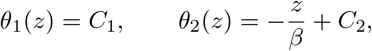

from which follows 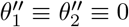. The last two terms of (30) satisfy (29) and so the second equation (30) becomes:

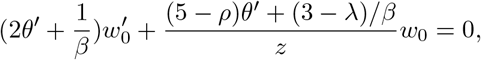

which has the following solutions: 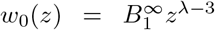 and 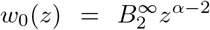, after we substitute *θ*_1_ and *θ*_2_, respectively. Then the leading term expansion of *w*(*z*) as *z* → ∞ we denote by *w*^∞^(*z*) (hence the superscript), which is following:

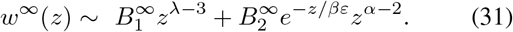

We perform the identical approach to obtain the asymptotic solution around zero. We let 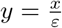 and (18) becomes:

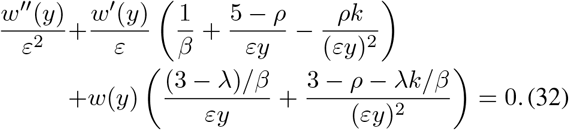

We are looking for the asymptotic expansion of the form (28). The functions *θ*(*y*) and *w*_0_(*y*) are now given by:

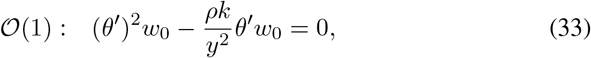

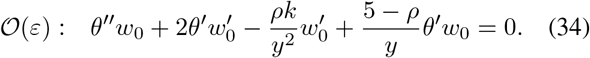

From (33) we obtain *θ*_1_ (*y*) = *C* and 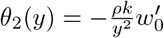, then the solutions of (34) are 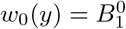 and 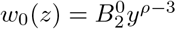 respectively. The leading order term expansion *w*^0^(*x*) of *w*(*x*) around zero is:

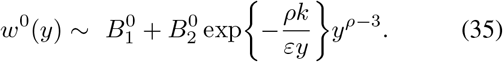

## Appendix III

### Algorithm for central connection problem

In this appendix, we describe the numerical procedure used to determine the dominant eigenvalue *λ* for (18). The asymptotic analysis in Appendix II provides the required behaviour of the solution near the singular points *x* → 0 and *x* → ∞. These conditions define a boundary-value problem on the interval (0, ∞), which is challenging to solve numerically. We therefore reformulate the problem as a central connection problem. Local solutions are obtained by solving initial-value problems derived from (18) that start close to the corresponding singular points and end at a matching point. The eigenvalue *λ* is then determined by requiring that these two solutions be linearly dependent, i.e., requiring that the Wronskian is equal to zero. The algorithm is as follows:

1. Rewrite initial equation (18) as a system of ODEs:

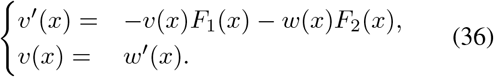
2. Set the value of parameters *α, β, k, λ*_max_, set *ε* = 0.01.
3. Split the domain 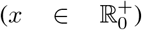 into two intervals 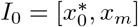 and 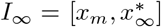, where *x*_*m*_ is a point of “matching”, where both solutions are significant, 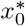 is a point close enough to zero, and 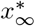 is a large value approximating infinity. The chosen approach is sensitive to values of *I*_0_ and *I*_∞_ bounds. To support numerical stability, we use simulations to determine these values. We empirically find that usually sufficient values are: the most frequent value of the sample density function based on simulation for *x*_*m*_, half of minimal reached concentration for 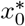 and maximal reached concentration for 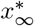.
4. Let (*w*_*L*_, *v*_*L*_) be solutions of (36) on *I*_0_, where the subscript ‘*L*’ denotes the solution to the left of *x*_*m*_. The respective initial conditions are given by the following vector 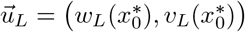:

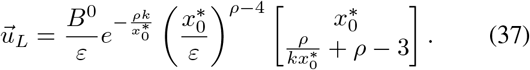 Similarly, let (*w*_*R*_, *v*_*R*_) be solutions of (36) on *I*_∞_ where the subscript ‘*R*’ denotes the solution to the right of *x*_*m*_. The respective initial conditions are given by the following vector 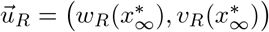:

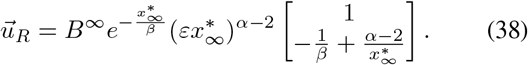 The initial conditions (37)–(38) are obtained from WKB approximations (35) and (31).
5. Let *f*(*λ*) be the Wronskian of functions *w*_*L*_ and *w*_*R*_ dependent on the parameter *λ*:

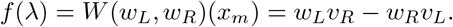

We require *w*_*L*_ and *w*_*R*_ to be linearly dependent at the matching point *x*_*m*_, which implies that the Wronskian vanishes. Thus, we use a root-finding method to determine values 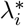 such that 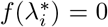 on the interval [0, *λ*_max_]. If no roots are found, then typically 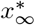 is insufficiently large or *ε* is chosen improperly.

6. For each given 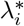, we compute the numerical solution of (18) *w*_num_ = (*w*_*L*_, *w*_*R*_) (Figs. 3A and 3C). The constants *B*^∞^ and *B*^0^ allow to arithmetically match *w*_*L*_ and *w*_*R*_, so that *w*_*S*_(*x*_*m*_) = *w*_*L*_(*x*_*m*_) and normalise solution to one for the next step.

7. We advert to requirement for the solution of (18) to be at least of class *C*^1^:

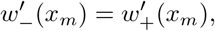

where

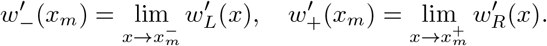

In case of multiple roots, *λ*_num_ is one with minimal error 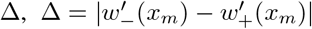. An example of smooth and non-smooth solutions are shown in Fig. 3B and 3D, respectively. In case of single root, if the solution is not smooth, then some roots of *f*(*λ*) are missing and numerical parameters need refinement. The derivative is obtained using local polynomial fit on both sides of *x*_*m*_ (instead of direct numerical derivative) due to its numerical robustness.

8. The solution of (15) for 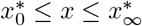 is given by:

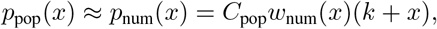

where *C*_pop_ is the normalisation constant determined through numerical integration of *w*_num_(*x*).

